# An Information-based approach to handle various types of uncertainty in Fuzzy Bodies of Evidence

**DOI:** 10.1101/706218

**Authors:** Atiye Sarabi-Jamab, Babak N. Araabi

**Affiliations:** School of Cognitive Sciences, Institute for Research in Fundamental Sciences (IPM), Tehran, Iran; Control and Intelligent Processing Center of Excellence, School of Electrical and Computer Engineering, University of Tehran, Tehran, Iran

**Keywords:** Distance of evidence, Fuzzy evidence theory, Fuzzy body of evidence, information-based comparison, set of most Discriminative Dissimilarity Measures (smDDM)

## Abstract

Fuzzy evidence theory or fuzzy Dempster-Shafer Theory captures all three types of uncertainty usually contained in a piece of information within one framework: fuzziness, non-specificity, and conflict. Therefore, it is known as one of the most promising approaches for practical applications. Quantifying the difference between two fuzzy bodies of evidence has a central role when this framework is used in applications. This work is motivated by the fact that while dissimilarity measures have been surveyed in the fields of evidence theory and fuzzy set theory, no comprehensive survey is yet available for fuzzy evidence theory. Here, we modified a set of the most discriminative dissimilarity measures (smDDM)-as the minimum set of dissimilarity with the maximal power of discrimination in evidence theory- to handle all types of uncertainty in fuzzy evidence theory. Consequently, our generalized smDDM (FsmDDM) together with one previously introduced fuzzy measure make up a set of measures that is comprehensive enough to compare all aspects of information conveyed by the fuzzy bodies of evidence. Experimental results are presented to show the efficiency of the proposed method.

## 1. INTRODUCTION

Dempster-Shafer theory (DST) or evidence theory is widely accepted as a flexible framework to model various processes of quantitative reasoning and decision making under uncertainty [1] [2] [3]. To make DST manage fuzzy information effectively in evidential reasoning, several generalization methods of DST to fuzzy sets are proposed [4] [5] [6] [7] [8] [9]. Computing a Fuzzy Body of Evidence (FBoE) can quantify different sort of uncertainties, such as imprecision, discord and degree of confidence.

Fuzzy DST is one of the most promising approaches for practical applications such as belief function approximation [10], regression analysis [11], risk analysis [12] [13] [14]. In all applications, measuring the difference between two FBoEs is a challenging task, as the dissimilarity measures have a central role in the body of the algorithms.

In fuzzy set theory, Bloch proposed a detailed survey of distances between fuzzy sets [15], where fuzzy distances were used in image processing applications. Also, de Campos et al. proposed a method to find distances between fuzzy measures based on associated probability distributions [16]. In DST, many works on measuring the distance between belief functions have emerged. In our previous study [17], we proposed a framework for comprehensive assessment of dissimilarity between two BoEs. Our outcome was the set of most discriminative dissimilarity measures (smDDM) that represents the minimal set of dissimilarity measures needed for an overall evaluation of the differences between two BoEs.

This paper is motivated by the fact that while dissimilarity measures in the field of evidence theory and fuzzy set theory have been studied separately, no comprehensive survey is yet available for fuzzy evidence theory to handle all types of uncertainty.

The rest of the paper is organized as follows: Section 2, gives a background on DST and fuzzy DST. The set of discriminative dissimilarity measures (smDDM) for DST is introduced in Section 3. The motivation of our work and our method to extend the smDDM to use in fuzzy evidence theory are proposed in Section 4. The interest of our suggestion is illustrated through experiment in Section 5. The paper is concluded in Section 6.

## 2. BACKGROUND

This section presents basic concepts in DST and fuzzy DST.

### 2.1 Dempster-Shafer Theory (DST)

DST or evidence theory assign mass values to the subsets of Frame of Discernment (FoD), instead of its elements. Let Θ be the FoD as a finite discrete set with N hypotheses, Θ ={ω_1_,…,ω_*N*_}, a mass function *m*(.), also called a Basic Probability Assignment (BPA) is defined as:

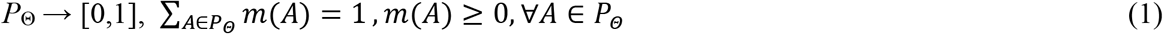

Here the *P*_Θ_ is the power set of Θ. Those subsets of Θ with non-zero mass values are called Focal Elements (FEs). All non-zero mass values (i.e. FEs along with their mass values) form a BoE. A BoE with *n* FEs is defined as:

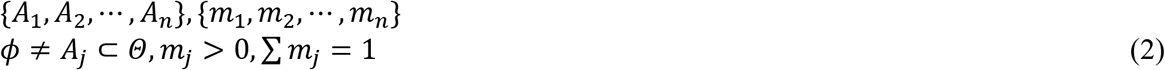

Given the mass functions of a BPA, a belief function Bel, and a plausibility function, Pl are introduced as [1] [2]:

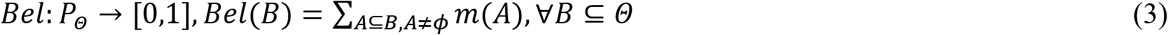

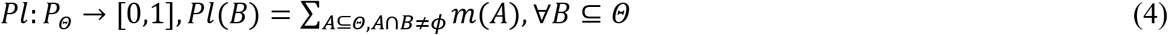

The combination of two independent bodies of evidence with the corresponding their mass functions *m*_1_(.) and *m*_2_(.) is given by the Dempster’s rule of combination as defined in [1].

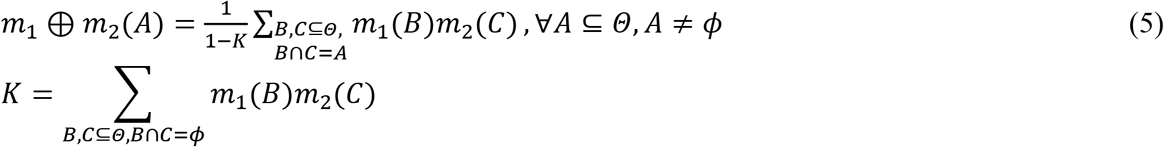

### 2.2 Fuzzy Dempster-Shafer Theory

The concept of fuzzy set has been proposed by Zadeh [18]. Since then, the analyses of fuzzy-valued data have become increasingly important [19]. With the preferred to manage imprecise and vague information in evidential reasoning, researchers tried to generalize the DST to deal with fuzzy sets [5] [7] [9] [20]. Fuzzy evidence theory extends DST to allow the assignment of degrees of belief to ambiguous propositions such as typically expressed in verbal statements, and represented by fuzzy subsets of the FoD.

A fuzzy body of evidence is defined as the following set

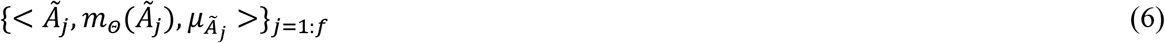

where Θ is a FoD and each 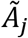 is a normal fuzzy set, such that 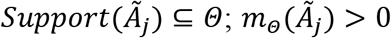, 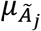 is the membership function of the fuzzy set 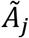;

Combining multiple fuzzy evidence structures can be extended through the Dempster’s rule by replacing the intersection of crisp sets - with the intersection of fuzzy sets. Following this idea, several rules were proposed to generalize the conjunction to fuzzy belief structure [21] [22]. Ishizuka et al. in [5] extended Dempster’s combination rule to fuzzy sets by considering the intersection degree of two fuzzy sets. Yen extended Dempster’s combination rule to fuzzy sets by using two operators with a cross-product operation and normalization process [22]. Yang et al. extended Dempster’s combination rule to fuzzy sets by constructing a weighted variable as follows [21]:

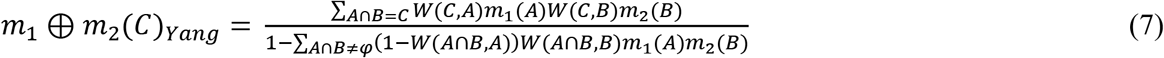

where, 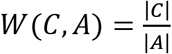 expresses the weight of contribution to the fuzzy set *C* from a FE *A*.

## 3. INFORMATION-BASED DISSIMILARITY ASSESSMENT IN DST

Dissimilarity assessment is a main problem in DST, where quantifying the difference between two BoEs has a central role in many practical algorithms. More than 60 dissimilarity measures have been used for DST in different applications. The idea of multi-dimension dissimilarity measures was first proposed by Liu [23] as a one-dimensional measure is inadequate to quantify the conflict between BoEs. Jousselme and Maupin reviewed and categorized the dissimilarity measures in DST, [24]. They focused on different formal properties of dissimilarity measures, selected 15 dissimilarity measures, and classified them into metric, pseudo-metric, semi-pseudo-metric and non-metric classes. However, the dissimilarity measures belong to each proposed class were highly correlated and they could not find one dissimilarity measure as a representor for each class. We developed a methodology to select a set of dissimilarity measures with the most discrimination power [17]. We investigated almost among all studied measures, used a forward selection procedure based on a proposed criterion to result in a set of measures that were maximally uncorrelated, while the discrimination was kept as high as possible. Our Outcome was the set of most Discriminative Dissimilarity Measures (smDDM) as:

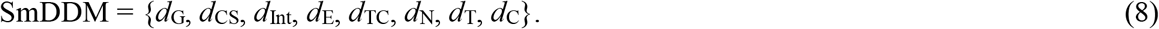

These members are classified into two classes: 1) intrinsic class, which relates to the imperfection of information provided by the sources of evidence ({*d*_*G*_, *d*_*E*_, *d*_*TC*_, *d*_*N*_, *d*_*C*_}), and 2) extrinsic class, which relates to the conflict/contradiction between BoEs ({*d*_*CS*_, *d*_*Int*_, *d*_*T*_}), where the dissimilarity are measured based on following criteria:

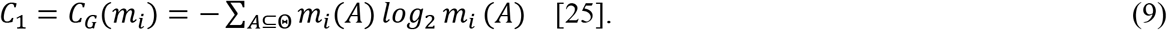

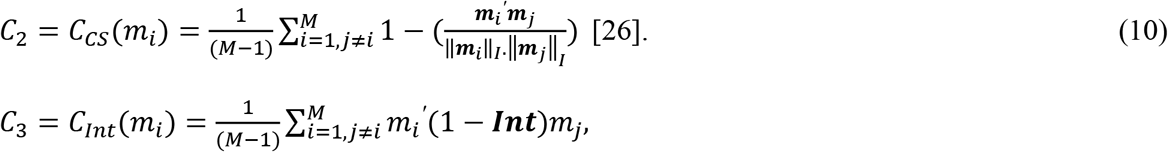

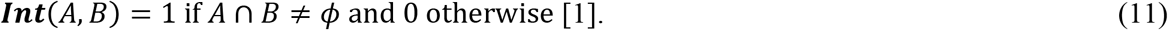

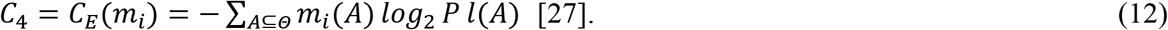

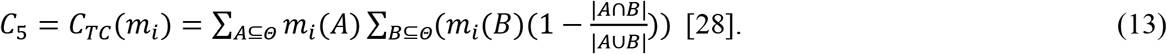

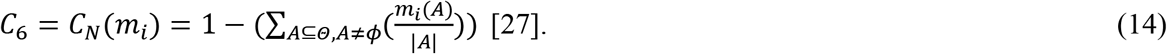

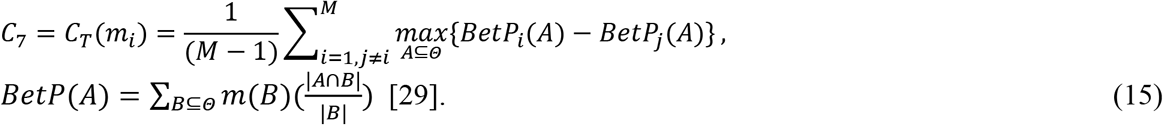

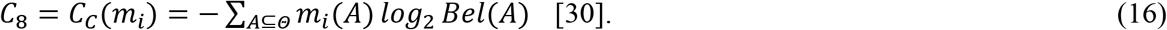

The results in [17], [10], and [13] showed that the smDDM can be an appropriate and justifiable disagreement measure for various applications. A more efficient description of the difference between two BoEs is obtained when these measures are considered together. However, they could just handle different types of non-specificity, and conflict, (i.e. strife, imprecision, and disparity). In following (Section 4), we extend the smDDM to compare all aspects of information conveyed by fuzzy bodies of evidence (i.e. fuzziness, non-specificity, and conflict).

## 4. THE MOTIVATION OF OUR WORK

The aim of our approach is to find a set of measures to handle different types of uncertainty between two FBoEs, i.e. fuzziness, non-specificity, and conflict. Although the smDDM can be an appropriate to handle non-specificity and conflict, the measuring of fuzziness was not included. Moreover, it could not use for fuzzy framework. In this section, first, we introduce the measures have been proposed in the fuzzy evidence framework (Section 4.1), then we extend the smDDM to apply for FBoEs (Section 4.2), and add these extended measures to the previously introduced ones to find a set could handle different types of uncertainty. However, among these criteria, some have redundant information content. To find the most important criteria, a backward elimination procedure is proposed to eliminate the lower significant criteria with the least salient properties (Section 4.3).

### 4.1. Uncertainty Measure in the Fuzzy Evidence Framework

For the fuzzy evidence framework, two main measures have been proposed [31]. The first measure is the General Uncertainty Measure (GM), which was introduced by Liu in his Ph.D. thesis [31].

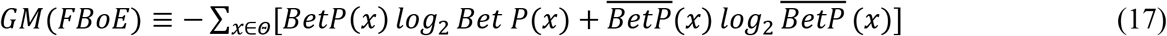

where,

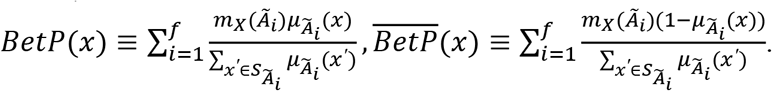

The second one is the Hybrid Entropy (FH), proposed by Zhu and Basir [32]. It was proposed as a measure which quantifies the overall uncertainty contained in a fuzzy evidence structure.

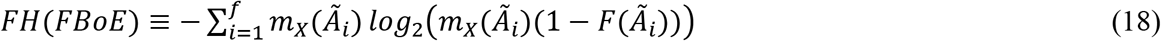

where 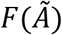 denotes the fuzzy entropy of fuzzy set 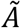 as:

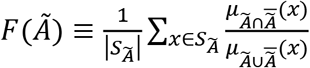

The smaller is 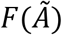, the less fuzzy is the fuzzy set 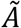.

These two measures are as the extension of the ambiguity measures (i.e. non-specificity and conflict) to fuzzy set. Therefore, basically they could just handle types of non-specificity, and conflict in fuzzy evidence theory.

Moreover, these measures were studied through a Monte Carlo simulation [31]. In this previous literature, results revealed that the GM has more stable behaviors, and the FH had counter-intuitive behavior in reflecting the changes in the conflict in FBoEs. Although both GM and FH claimed that quantified aggregate uncertainty is an extension of the ambiguity measure (non-specificity and conflict) to fuzzy set, the differentiation of the quantity of the three types of uncertainty, i.e. fuzziness, non-specificity, and discord is not possible.

There are also some criteria which try to measure fuzziness in an FBoE. A brief summary about the fuzziness measures of fuzzy sets had been introduced in [33]. The fuzziness of an FBoE is estimated as a weighted sum of fuzziness over all different FEs of FBoE as follows:

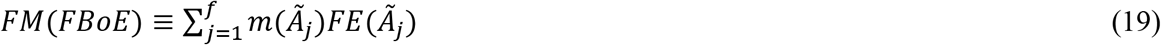

where the 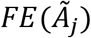 is a function to measure the fuzziness of each fuzzy set, which can be measured by using the quantity proposed by De Luca and Termini [20].

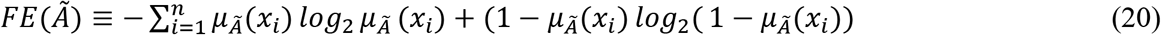

This measure was corresponding to Shannon’s probabilistic entropy. The extended version can be found in [34], [35], [36] and [37].

### 4.2. Generalizing the smDDM to Fuzzy DST (FsmDDM)

To extend the smDDM for an FBoE, the FBoE should be replaced by BoE. A natural way to link a piece of fuzzy evidence with crisp evidence is to represent the fuzzy set by its α-cuts using the resolution identity principle [26], and then distribute 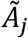’s mass 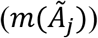 to its α-cuts. This process can be described as follows:

1. *Decompose the fuzzy FE to its α-cuts*. For each α ∈ [0,1],

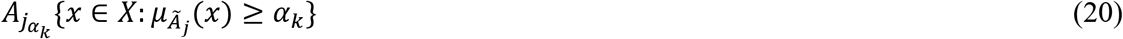
2. *Distribute the mass of fuzzy FE to its α-cuts*.

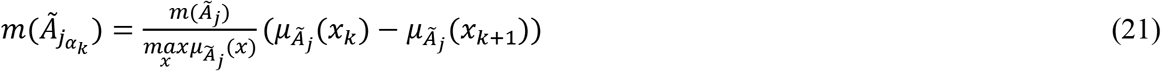

which ensures that 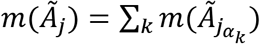

Therefore, we can calculate dissimilarity measures using smDDM in classical DST. Then we will have a collection of dissimilarity measures for FsmDDM. We could use the averaging of these measures, and finally have a FsmDDM which measure all aspects of information between two FBoEs.

### 4.3. Finding the most important criteria

Up to now, we separately estimated the ambiguity and fuzziness with a given FBoE. We add these extended measures to the previously introduced ones (i.e. GM in Eq. (17), and FH in Eq. (18)) to find a more riche set of measures. Moreover, the addition of the fuzziness measure (i.e. FM in Eq. (19)), results a vector which could better handle different types of uncertainty. However, among these criteria, some may have redundant information content.

To find the most important criteria, a backward elimination procedure is proposed to eliminate the lower significant criterion. The advantage of backward elimination is that it gives the opportunity to look at all criteria before removing the least salient one. Finally, we opt a comprehensive set of criteria.

To apply backward elimination, a sample of possible values used on all criteria should be selected. In the first step, 1000 random FBoEs is generated by adapted algorithm which is proposed in [31] based on Tessem’s idea [29]. Algorithm I summarize the generating procedure.

**ALGORITHM I.**
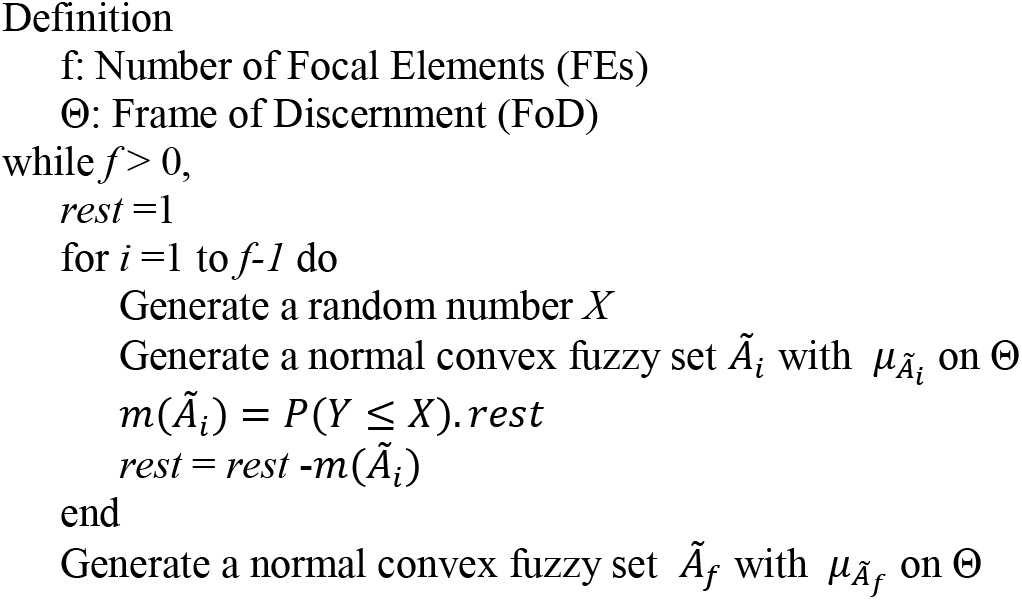
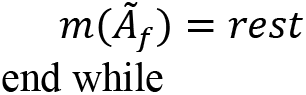
GENERATINGING A RANDOM FBOE.

Then, we assess each FBoE based on all criteria (i.e. FsmDDM, GM, FH, and FM). As a result, a source value is obtained. The backward elimination starts with 4 criteria and removes the criterion with least information in each step as follows:

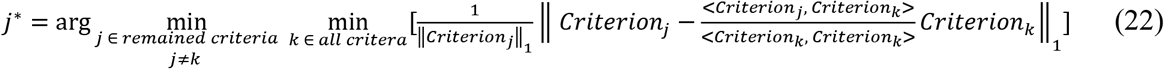

In each step of backward elimination, the normalized distance between the *j*th criterion and its orthogonal projection on the *k*th criterion, since the *j*th criterion has more independent information content related to other criteria, the scoring criterion increases. In the worst case, the minimum of this scoring criterion for all criteria is taken in J(*j*).

Fig. 1 shows the score of selection in order of removal, where 4 criteria (i.e. FsmDDM, GM, FH, and FM) are considered. After removing FH, and GM, the graph increases. This means that choosing the first two removed criteria does not convey much more information. As a result, the FsmDDM along with the fuzzy measure (i.e. FM) result a vector which could better handle different types of uncertainty.

**Fig. 1.**
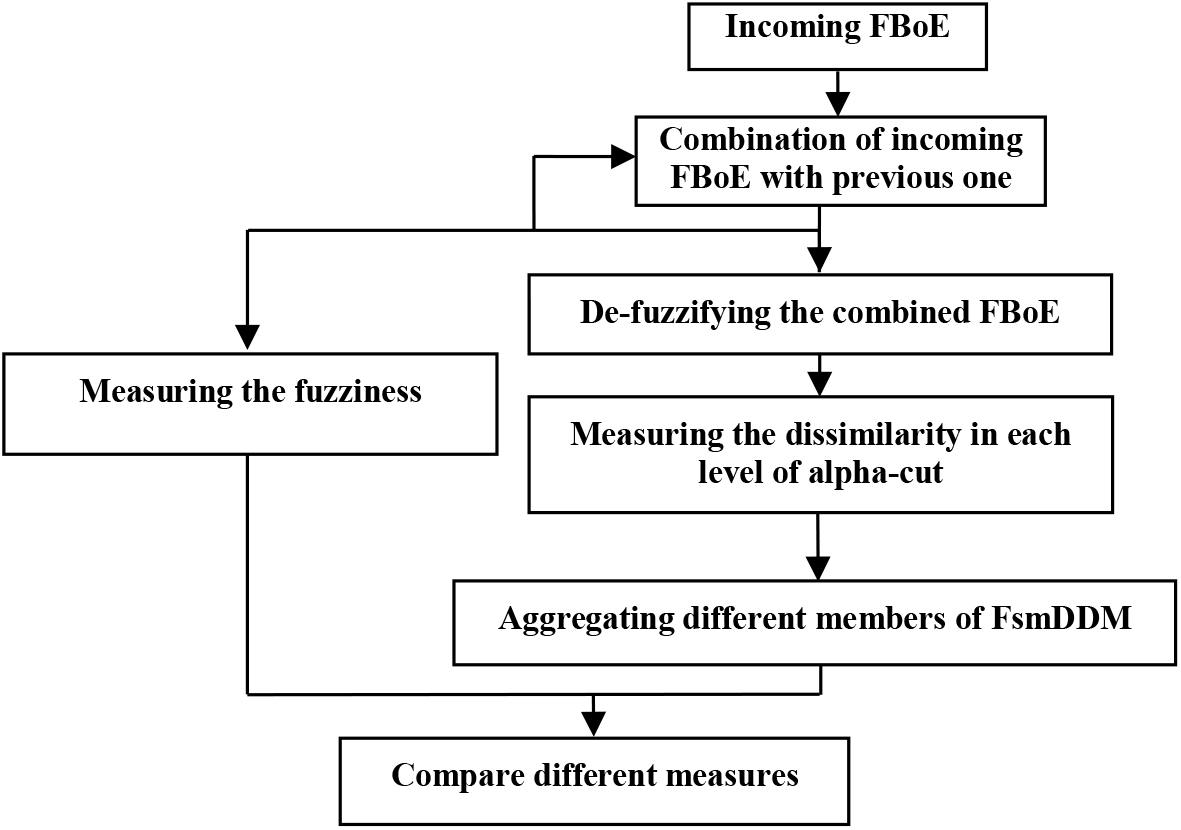
Score of selection. Removing the two criteria yields not much unimportant criteria.

## 5. EXPERIMENTAL RESULTS FOR EVALUATING THE FSMDDM

In this section, we evaluate the FsmDDM as well as FH, and GH. As these measures are not enough used in real application in literature, we compare the accuracy and reliability of proposed measure through simulation.

### 5.1. Difference between measures through simple examples

In order to examine the behavior of the dissimilarity measures in detail, we first generate simple numerical examples FBoEs, allowing us to predict the changing of the content of information and to compare the values given by different measures.

Assuming three fuzzy bodies of evidence as follows:

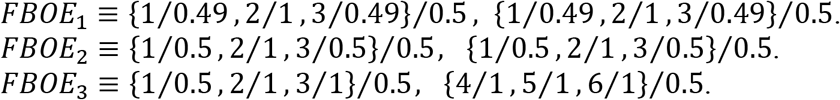

In this example, one can see that the fuzziness is a little smaller in *FBOE*_1_, and remains the same for the two others. The discord, however, almost the same for *FBOE*_1_ and *FBOE*_2_ and it changes dramatically from *FBOE*_2_ and *FBOE*_3_. Here are the values of measures for these BoEs.

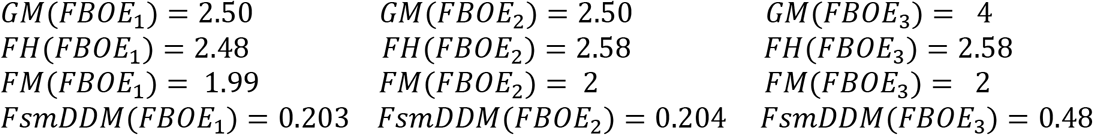

The results show that the FsmDDM could be better in reflecting changes in all types of information.

### 5.2. Difference between measures through average behavior during combination of EBoEs

We use a Monte Carlo simulation as is done in similar works [31]. The Monte Carlo simulation is used in a combination procedure. The process of combining can reflect the uncertainty decrease when the multiple FBoEs are combined sequentially. Fig. 2 shows the procedure of combination process and information-based comparison in general.

**Fig. 2.**
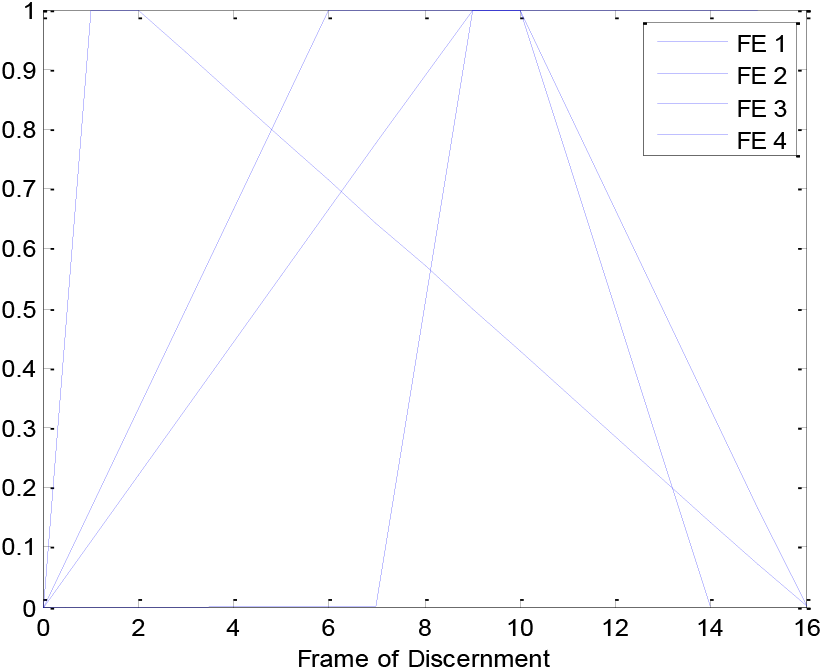
Information-based comparison approach during combination process between two fuzzy bodies of evidence.

A random FBoE is generated by adapted algorithm which is proposed in Algorithm I. In our experiment, |*Θ*| = 16, four fuzzy FEs for each new incoming FBoE are considered. Fig. 3 illustrates one FBoE with four normal trapezoidal fuzzy numbers as its FEs, which is randomly generated through Algorithm I.

**Fig. 3.**
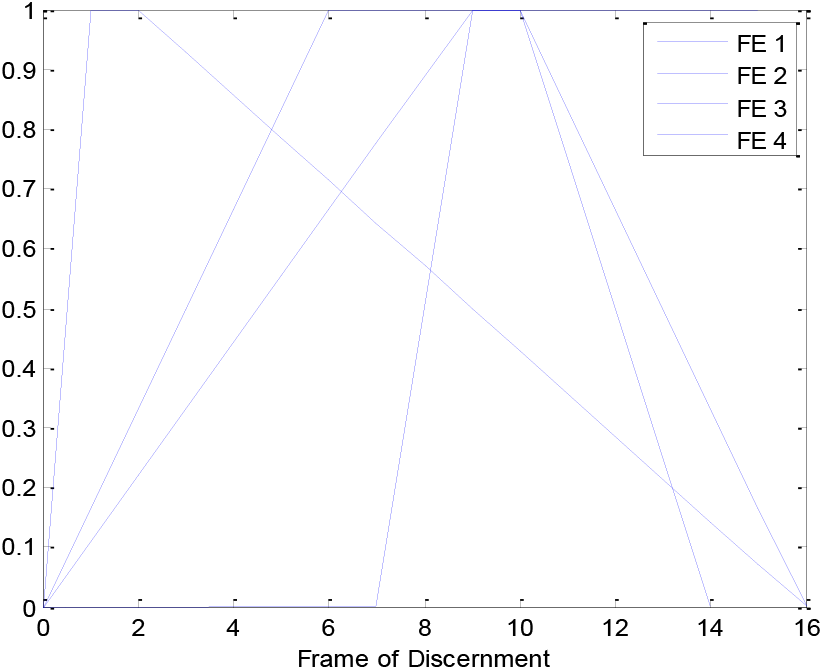
A FBoE with four normal trapezoidal fuzzy focal elements.

During the experiment, 20 combinations are assumed. In order to combine the information provided by each piece of evidence in fuzzy DST, we use the Yang et al. combination’ rule as Eq. (7). To compute the FsmDDM, the new combined FBoE has to be normalized fuzzy set. Here, we used the normalization method proposed by Yager and Filev [38]. Their method is based on the scaling up. The normalization process is presented in Algorithm II.

**ALGORITHM II.**
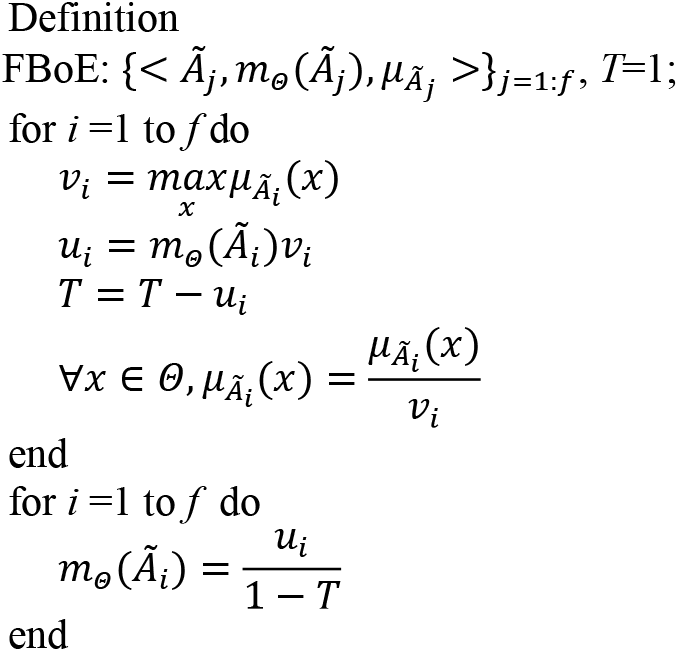
NORMALIZATION PROCESS.

In each combination step, the difference between combined FBoE with the previous FBoE is measured as follows:

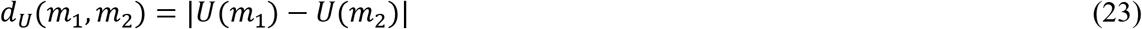

where, *U* can be any measure. Then, the experiment is repeated 50 times and the mean of differences is calculated. Fig. 4 shows the resulting curves. The results are in line with the results reported in [31], here, although GM is more stable that FH, the behavior between two consecutive combination steps does not decrease all the time. As shown in Fig. 4, our proposed overall dissimilarity based on FsmDDM has more stable behavior between two consecutive combination steps in the sense that the average curve of our proposed dissimilarity is more stable than GM and FH. However, the aggregate uncertainty measures of GM and FH have no acceptable behavior during combination.

**Fig. 4.**
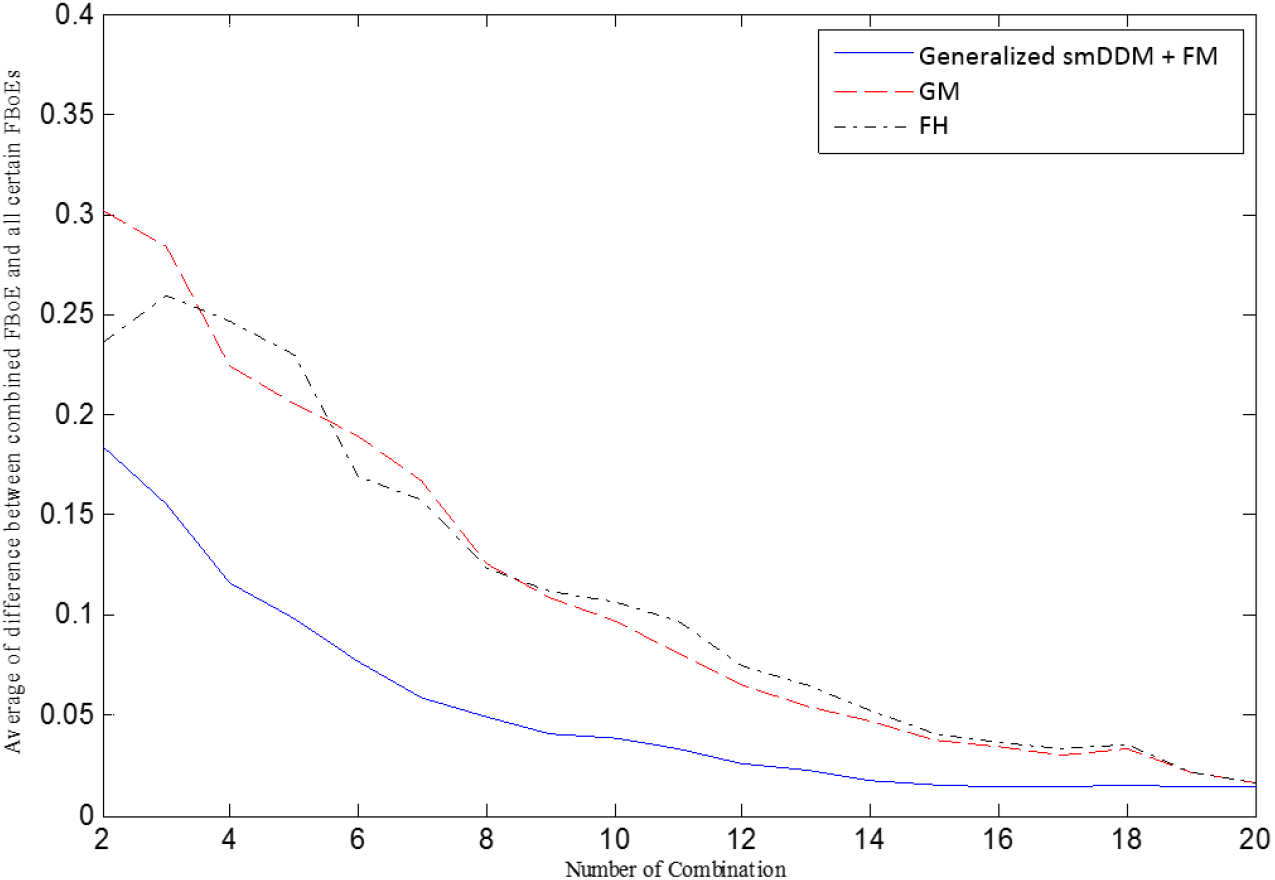
Mean of dissimilarity measures, when the combined FBoE is compared with all certain FBoEs, here the dash line shows the mean of FsmDDM plus fuzziness measure.

## 6. CONCLUSION

Fuzzy evidence theory captures all three types of uncertainty usually contained in a piece of information within one framework: fuzziness, non-specificity, and conflict. Quantifying the difference between two FBoEs has a central role when this framework is used in applications.

In this paper, we found a method to compare two FBoEs comprehensively.

Although our previously proposed smDDM could be an appropriate to handle non-specificity and conflict, the measuring of fuzziness was not included. Moreover, the smDDM could not use for fuzzy framework. In this study, we extended the smDDM to apply for FBoEs, and tried to add it to the previously introduced ones to find a rich set of measures which could handle different types of uncertainty. However, among these criteria, some have redundant information content. We proposed a backward elimination to find the most important criteria. In the proposed backward selection procedure, a scoring criterion was designed to remove unimportant DMs gradually. Finally, we selected a set of most important ones and showed why the selected criteria were efficient. Consequently, we had a set of dissimilarity measure which can measure all three types of uncertainty: fuzziness, non-specificity, and conflict.

We investigated the ability and stability of our proposed measure through a Monte-Carlo simulation. The results showed that the trend curves have more acceptable behavior when the FsmDDM is used.

## CONFLICTS OF INTEREST STATEMENT

The authors have declared that no competing interests exist.

